# Rational design of novel fluorescent enzyme biosensors for direct detection of strigolactones

**DOI:** 10.1101/2020.03.10.986562

**Authors:** Rebecca J Chesterfield, Jason H Whitfield, Benjamin Pouvreau, Da Cao, Christine A Beveridge, Claudia E Vickers

## Abstract

Strigolactones are plant hormones and rhizosphere signalling molecules with key roles in plant development, mycorrhizal fungal symbioses, and plant parasitism. Currently, sensitive, specific, and high-throughput methods of detecting strigolactones are limited. Here, we developed genetically encoded fluorescent strigolactone biosensors based on the strigolactone receptors DAD2 from Petunia hybrida, and HTL7 from Striga hermonthica via domain insertion of circularly permuted GFP. The DAD2 biosensor exhibited loss of cpGFP fluorescence in vitro upon treatment with the strigolactones 5-deoxystrigol and orobanchol, or the strigolactone analogue GR24. The biosensor likewise responded to strigolactones in an in vivo protoplast system, and retained strigolactone hydrolysis activity. The ShHTL7 biosensor exhibited loss of cpGFP fluorescence upon GR24 treatment in vitro, and responded to a specific inhibitor of ShHTL7 but not DAD2, indicating that the biosensors retained the specificity of their parent receptors. These biosensors have applications in high-throughput screening, and may also have utility for studying strigolactone biology.

## Introduction

Strigolactones (SLs) are plant hormones involved in nutrient distribution, environmental adaptation, and plant development. They also function as rhizosphere signalling molecules in both symbiotic and parasitic interactions. Hormonal functions of strigolactones include branching inhibition, regulating root development and root hair growth, inhibiting adventitious rooting, increasing secondary growth, and promoting leaf senescence ^1–6^. SLs have also been shown to improve plant survival and productivity under drought and salinity stress ^7–9^. As rhizosphere signalling molecules, SLs induce hyphal branching in arbuscular mycorrhizae, promoting symbioses that are important for nutrient access in over 80% of land plant species ^10^. However, SLs also trigger seed germination of *Striga* and *Orobanche spp*. of parasitic plants, and therefore play a major role in parasitisation of crop plants ^11^. These parasitic species infest upwards of 60 million hectares of farmland worldwide, parasitising most major cereal crops and leading to major economic burden (losses reach $US billions annually) and food insecurity in endemic regions ^12^.

The diverse functions of SLs (and SL-like compounds) lend themselves well to agricultural applications. A number of SL analogues have been investigated as *Striga* and *Orobanche* control agents (by inducing suicidal seed germination) with promising results ^13–15^. SLs may also act as biofertilisers by enhancing mycorrhization, and their regulation of drought and salinity responses may provide the capacity to improve crop productivity under unfavourable conditions ^16^. Furthermore, use of SLs to control plant architecture may contribute to the development of high-density, high-productivity planting systems, while crops with modified strigolactone production, composition, or perception show potential for improved agronomic properties ^17–20^.

The biology and physiology of strigolactones is not yet fully elucidated, and there are indications that novel unidentified strigolactones exist ^21^. A high-throughput detection method would facilitate their identification by allowing screening of a wide range of plant tissues and extracts. High-throughput detection would also simplify chemical library screening to identify strigolactone mimics with desirable agricultural properties, and accelerate metabolic engineering in microorganisms for development of strigolactone production platforms, which are of particular interest due to the difficulty of chemical synthesis ^16^.

Currently, no cost-effective, high-throughput methods for specific SL detection exist. Strigolactones are typically detected and measured by mass spectrometry, requiring expensive equipment and analytical standards. Functional assays, including parasitic seed germination and/or branching inhibition studies offer an alternative method of SL detection; however, accidental release of parasitic seeds is an environmental risk, and branching assays can be labour intensive. Molecular biosensors offer an alternative to existing assays with the potential for development as high-throughput systems.

*In vivo* SL biosensors have previously been developed by adapting the endogenous perception machinery. SLs are bound by the SL receptor D14 (known as DAD2 in *P. hybrida*), triggering a conformational change that facilitates interaction with the SCF E3 ubiquitin ligase F-box protein MAX2 (known as D3 in rice (*Oryza sativa*)), and the transcriptional repressor D53 (or SMXL6/7/8 orthologues in *Arabidopsis thaliana*) (Figure 1). This interaction triggers ubiquitination and degradation of D53, mediating downstream transcriptional changes ^22^. In addition to signal transduction, D14/DAD2 hydrolyses SLs, potentially as a feedback mechanism to downregulate SL signalling ^23^. As an additional feedback mechanism, SL signalling also induces degradation of D14/DAD2 ^24^ (Figure 1). D14 degradation is dependent on MAX2 and coupled to D53 degradation; however it remains unknown whether D14 is a direct ubiquitination target of the SL-induced SCF complex, or is the target of a ubiquitin ligase expressed in response to D53 degradation ^25^.

**Figure 1.**
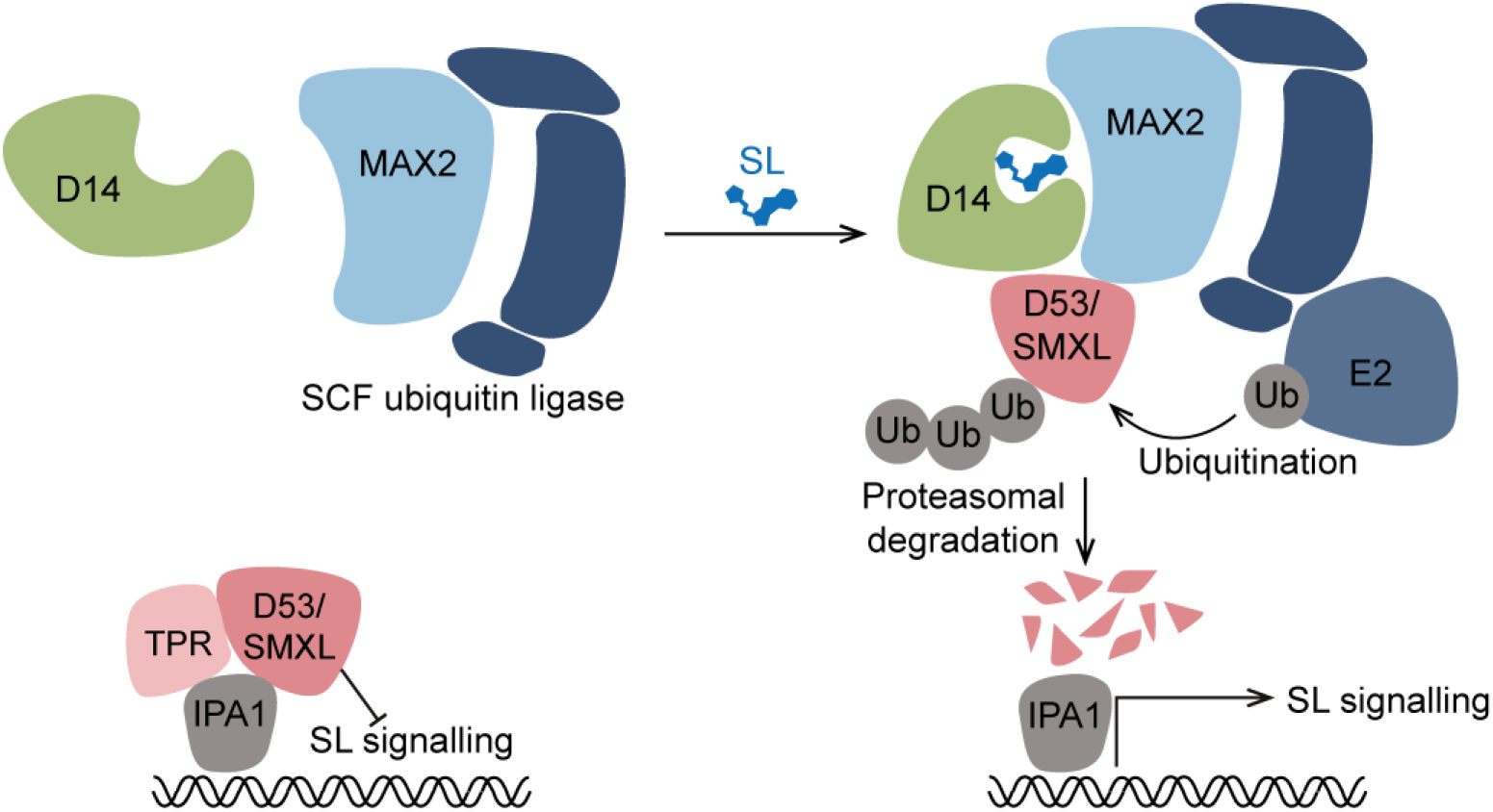
Strigolactone perception machinery. Schematic illustrating strigolactone signal propagation in plants. In the absence of SL, D53 (or SMXL6/7/8 orthologues) repress downstream SL signalling in complex with SPL transcription factor IPA1, and TPR corepressors. SL perception by D14/DAD2 triggers interaction with D53 and MAX2, the substrate-recognition component of an SCF ubiquitin ligase complex. The SCF ligase ubiquitinates D53, targeting it for proteasomal degradation and causing transcriptional de-repression of SL signalling genes. SL signalling also results in D14/DAD2 degradation via an unknown mechanism.

Making use of the SL-induced degradation of D53, Samodelov *et al* ^26^ generated StrigoQuant, a D53-luciferase fusion that could detect SL signalling in *Arabidopsis* protoplasts. Similarly, Sanchez *et al* generated transgenic *A. thaliana* lines expressing a D14-luciferase fusion that could detect exogenously applied SLs and SL mimics by taking advantage of the endogenous feedback loop where SLs trigger degradation of D14 ^27^. Additionally, the SL-dependent interaction between DAD2 and MAX2 was initially demonstrated using a yeast two-hybrid (Y2H) system ^28^. A similar Y2H approach enabled a high-throughput yeast screen for parasitic seed germination stimulants that relied on the interaction between MAX2 with HTL, a paralogue of *Arabidopsis* D14 that is involved in the perception of karrikins and cotylimides ^29^. An analagous high-throughput yeast screen could conceivably be developed for SLs.

While the above multi-component SL biosensors provided new methods for strigolactone research, the outputs of such systems can be difficult to interpret due to the indirect nature of measurement as well as the lag time between signal input and observed signal output. Moreover, given the complexity of such systems requiring integration of multiple signalling events, modulation of individual components can introduce noise or signalling artefacts thus reducing confidence in the data. Direct detection of the target molecule would be ideal, however, design and construction of such biosensors is not straightforward owing to a lack of employable general design principles ^30^.

In recent years, several methods for the development of integrated fluorescent sensors have emerged, aiming to allosterically link the fluorescent intensity of a circularly permuted green fluorescent protein (cpGFP) variant to the activity or conformational state of a host receptor ^31–34^. Circular permutation interchanges amino and carboxyl fragments of a protein, reconnecting them via a short linker. This renders GFP highly sensitive to local conformational changes ^35^. The conformational sensitivity of cpGFP can be exploited in biosensor design by inserting it into a molecular receptor with large-scale ligand-dependent conformational transitions. While highly successful, these approaches have relied on random library generation and bespoke assay development, limiting the wider implementation of each method.

In this work, we sought to exploit the availability of molecular structures and biophysical characterisations of the DAD2 and ShHTL7 enzymes to rationally design domain insertion biosensors for SL detection. To our knowledge, this is the first instance in which this GFP insertion strategy has been applied to an enzyme to monitor its substrate. This sets a strong precedent for future biosensors whereby if ligand-dependent conformational changes occur in a given structure there is potential to create a fluorescent probe capable of direct detection of a target.

## Results

### Generation of an SL biosensor by FP domain insertion into DAD2

To develop a single-component SL biosensor, we applied the approach of fluorescent protein domain insertion into the *P. hybrida* SL receptor, DAD2. The fluorescent properties of circularly permuted fluorescent proteins are highly sensitive to local conformational changes ^35^, a property that has been exploited to develop several small molecule biosensors by inserting the circularly permuted fluorescent protein into an appropriate analyte binding protein ^31–33^. We therefore aimed to insert cpGFP into DAD2 in such a way that would preserve SL binding and propagate the DAD2 conformational change into the fluorescent protein domain, thus modulating fluorescence in a SL-dependent manner (Figure 2A).

**Figure 2.**
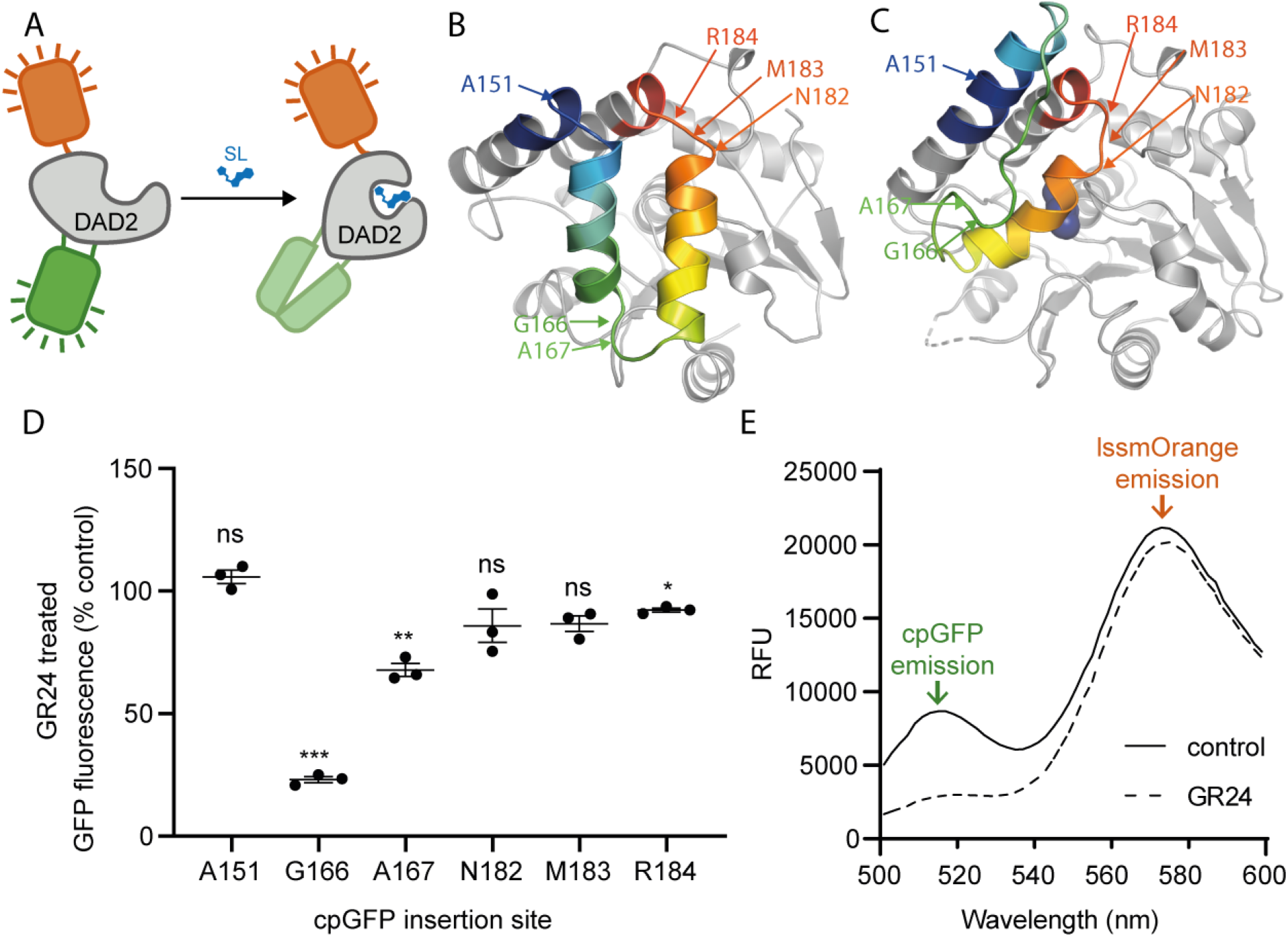
DAD2-based strigolactone biosensor using cpGFP loop insertion. (A) Schematic of the ratiometric DAD2 cpGFP biosensor design. Strigolactone-induced conformational change is propagated into cpGFP, reducing fluorescence. The C-terminal LSSmOrange fusion provides an internal fluorescent control. (B) Structure of D14 in its apo conformation, and (C) bound to a strigolactone hydrolysis product (navy spheres). A region of the lid domain that undergoes a large conformational change is highlighted in rainbow mode. The six cpGFP insertion sites are indicated with arrows on each structure (DAD2 numbering), where the indicated residue is immediately N-terminal of the cpGFP insertion. (D) Fluorescent response to the strigolactone analogue *rac*-GR24 at each insertion site. Error bars represent standard error of the mean. *p<0.05, **p<0.01, ***p<0.001 by one sample t-test compared to theoretical mean of 100 (no difference). (E) Emission spectrum of rDAD2cpGFP(G166).

The crystal structures of the *Arabidopsis thaliana* orthologue, *At*D14, (PDB ID 4IH4 – Apo ^36^ and 5HZG – Holo ^37^) were used to assess the scale and locality of ligand-induced conformational transitions. D14/DAD2 is composed of two major domains denoted here as the catalytic domain and lid domain, the latter of which comprises α helices 5-8 (residues 136 – 194; for clarity DAD2 numbering and secondary structure annotation are used throughout; see ^28^). Based on alignment of these structures using the catalytic domain, the DAD2 protein undergoes what appears to be a ‘lid-closing’ conformational change upon interaction with a target SL (Figure 2B, C). A key parameter for domain insertion sensors is insertion into flexible loop regions as these are often the most significantly transposed by any transitions but also the most tolerant of a domain insertion. Following these parameters and based on the structural alignment we focussed on the three loops between helices 5-8, that displayed the most significant transitions (Figure 2B, C). A set of six sites were selected for testing: A151, G166, A167, N182, M183 and R184. Based on previous studies of domain insertion sensors, a minimal linker of Ala-Ser was used to insert cpGFP into the target sites, as this appears sufficient to transduce minor loop rotations and perturbations in the host receptor ^31^.

Each domain insertion variant was expressed in *Escherichia coli* and cpGFP fluorescence was measured in lysates in the presence or absence of the SL analogue *rac*-GR24. CpGFP insertion at two sites, G166 and A167 (both located between helices 6 and 7) exhibited reduced fluorescence in the presence of *rac*-GR24 with insertion after G166 (denoted DAD2cpGFP(G166)) giving a four-fold reduction in fluorescence (Figure 2D, Supplementary table 1). In contrast, insertions in the loops between helices 5 and 6, or helices 7 and 8, gave low or no change in fluorescence in response to *rac-*GR24. It is possible that insertion in these loops did not result in the propagation of conformational changes into cpGFP, or alternatively that the insertion perturbed DAD2 structure, disrupting interaction with *rac*-GR24. Nevertheless, the signal changes arising from the G166 and A167 insertion sites validated the use of domain insertion biosensors for SL detection. DAD2cpGFP(G166) was pursued for further development.

Intensiometric sensors that utilise only a single-fluorophore, while effective as probes, have some limitations. In particular, intensiometric sensors are sensitive to changes in expression level and protein degradation, which are sources of significant noise for signal intensity. To rectify this, we sought to convert the biosensor to a ratiometric output independent of sensor concentration by adding a second fluorophore to provide an internal normalisation control (Figure 2A). LSSmOrange has a <u>l</u>ong <u>S</u>tokes <u>s</u>hift, allowing the cpGFP and LSSmOrange fluorophores to be excited by a single wavelength, while still providing spectrally resolved emission, as employed by the Matryoshka gCAMP sensors ^38^. This approach exploits the non-overlapping excitation-emission spectra of the fluorophores, eliminating confounding effects of FRET. LSSmOrange was fused at the C-terminus of the DAD2cpGFP(G166) via a rigid GEAAAKEAAAKGG linker (denoted here as rDAD2cpGFP(G166)). Treatment with SL showed retention of the cpGFP intensity switching; and importantly, LSSmOrange fluorescence intensity was unaffected (Figure 2E), enabling calculation of a dose-dependent ratio change.

### The DAD2 conformational biosensor has high sensitivity *in vitro*

To test the sensitivity and detection range of rDAD2cpGFP(G166) we initially measured the fluorescence ratio cpGFP/LSSmOrange 10 minutes after adding various concentrations of *rac*-GR24. *rac*-GR24 had a detection range approximately between 50 nM and 500 nM, with an apparent EC50 of 143 ± 16 nM (Figure 3A).

**Figure 3.**
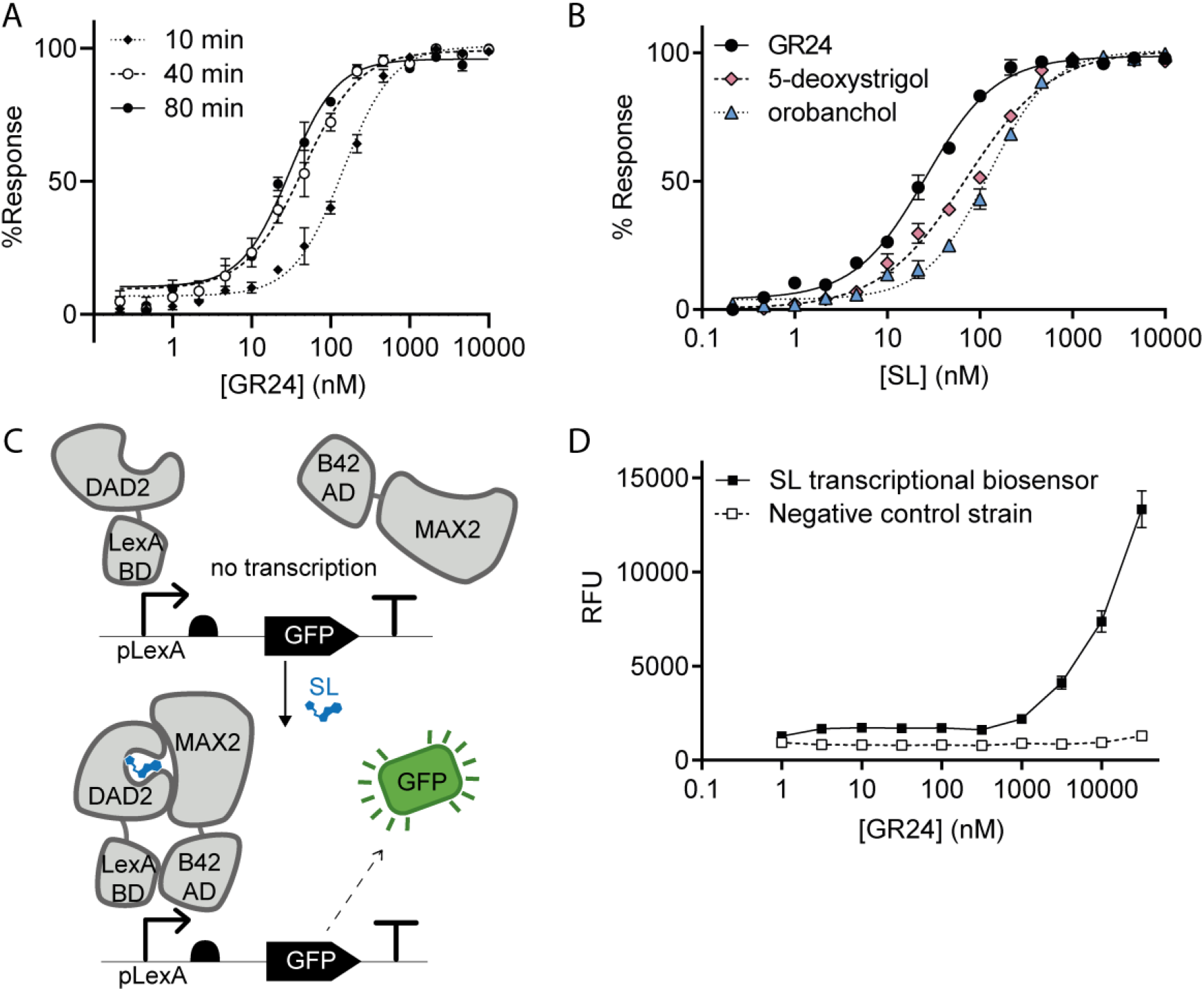
Sensitivity and detection range of rDAD2cpGFP(G166) compared to a two-component transcription-based biosensor. (A) Dose-response curves of rDAD2cpGFP(G166) after different incubation times. (B) Dose-response curves of rDAD2cpGFP(G166) to different SLs after 50 min incubation. (C) Schematic of the two-component transcription-based reporter. DAD2 is fused to the LexA DNA binding domain, and MAX2 is fused to the B42 activation domain. Strigolactones trigger interaction between DAD2 and MAX2, recruiting the activation domain to the promoter, and inducing transcription of the GFP reporter protein. (D) Dose-response curve of the transcriptional biosensor after 16 h incubation with *rac*-GR24. The transcriptional biosensor has lower sensitivity than rDAD2cpGFP(G166) and had not saturated the response at the highest *rac*-GR24 concentration tested. Data show mean ± standard error of three replicates.

Unlike existing domain insertion biosensors, DAD2cpGFP(G166) is not only a receptor, but also an enzyme. The relationship between SL hydrolysis and DAD2 conformational change is uncertain ^39^, and some but not all reports suggest that SL receptors form stable interactions with a reaction product ^23,37,40,41^. We hypothesised that the dose-response behaviour may be time-dependent as a result of the non-equilibrium system. Indeed, sensitivity and dynamic range were dependent on the incubation time, with the sensitivity of the biosensor increasing with incubation time (Figure 3A). This is consistent with the formation of a stable receptor-product complex locking the biosensor in a reduced fluorescence state.

In *vitro*, this time-dependent sensitivity may allow a larger dynamic range to be accessed simply by measuring at a range of time points. Additionally, sensitivity remained relatively stable between 40 and 80 min, which permits a large number of samples to be processed and measured in parallel without significant influence from incubation time. In our system, we could detect *rac*-GR24 concentrations down to low nanomolar concentrations using a 40 – 80 min incubation time, with an apparent EC50 of 29.9 ± 4.7 nM at 80 min (Figure 3A).

Next, we investigated the response of the biosensor to the natural SLs 5-deoxystrigol and orobanchol, choosing 50 min incubation to ensure stable readings. The biosensor responded in a dose-responsive manner to both natural strigolactones tested. The EC50 values were fit at 70.5 ± 6.3 nM for 5-deoxystrigol, and 124 ± 6.8 nM for orobanchol, which were significantly higher than that for *rac*-GR24 (EC50 = 25.9 ± 1.7 nM, extra sum-of-squares F test, p<0.0001).

### An alternate biosensor design had lower sensitivity

In parallel to rDAD2cpGFP(G166), we also developed a yeast two-hybrid (Y2H)-based reporter of SL as an alternative biosensor design. The Y2H reporter was similar to a design reported by Hamiaux *et al.* ^28^, but with different activation and binding domains. Transcriptional reporters are commonly used in biosensor development, especially in metabolic engineering applications. By fusing DAD2 to a DNA binding domain and MAX2 to an activation domain, transcription of the reporter gene (yEGFP) in *Saccharomyces cerevisiae* was coupled to SL-dependent interaction between DAD2 and MAX2 (Figure 3B). After 16 h incubation with *rac*-GR24, the Y2H approach had much lower sensitivity than the conformational biosensor (Figure 3C), potentially due to poor uptake of *rac-*GR24 by the cells, or alternatively due to minimum concentration requirements of the reconstituted transcription factor. Similarly, while the Y2H reporter developed by Hamiaux *et al.* had higher sensitivity than that developed here, the EC50 was approximately 10 times higher than that observed for the direct-detection domain insertion biosensor ^28^. As the Y2H approach had lower sensitivity, slower response times and was restricted to use in a single organism, the domain insertion biosensor was pursued for further application.

### The DAD2 conformational biosensor detects SL in protoplasts

Plant protoplasts can be used as a cell-based model for plant systems, providing a simple system for transient expression of proteins of interest. Accordingly, protoplasts have been used extensively in studies of strigolactone signalling ^25,26,42–44^. We therefore chose protoplasts as a model system for proof of concept of the DAD2 conformational biosensor function in plant cells.

Tobacco protoplasts transfected with rDAD2cpGFP(G166) were examined by flow cytometry, and exhibited fluorescence in GFP and LSSmOrange channels. Upon treatment with *rac*-GR24, GFP fluorescence decreased to just above background, whereas LSSmOrange fluorescence decreased only slightly, likely due to loss of bleed-through from cpGFP (Figure 4A). As observed in bacterial lysates, there was a slight time-dependent response: the apparent EC50 was 173.0 nM after 30 min, and decreased to 101.9 nM after 60 min (Figure 4B).

**Figure 4.**
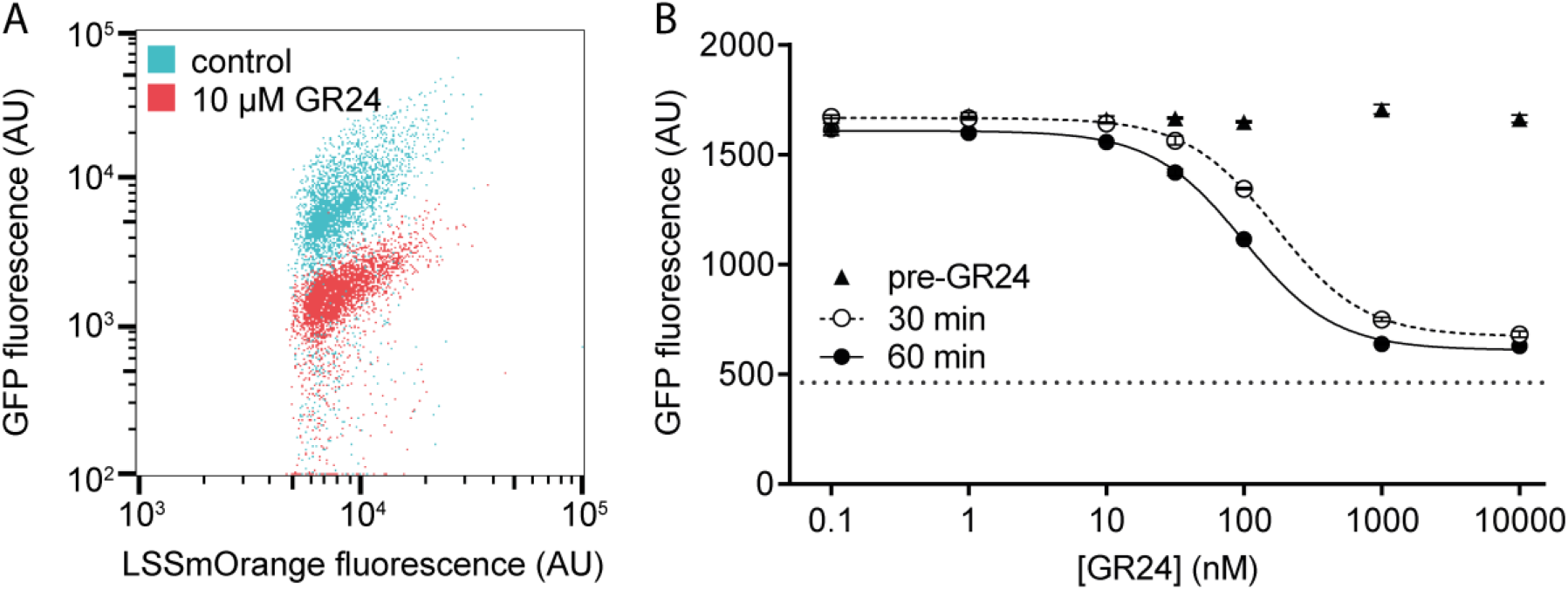
rDAD2cpGFP(G166) responds to *rac*-GR24 in protoplasts. (A) Dot plot of protoplasts expressing rDAD2cpGFP(G166) treated with 10 μM *rac*-GR24 or acetone control. For clarity, only cells gated for positive LSSmOrange fluorescence are shown. (B) Dose-response curve of protoplasts transformed with rDAD2cpGFP(G166) to *rac*-GR24. Each sample was measured prior to, 30 min after, and 60 min after addition of *rac*-GR24, and data represent mean fluorescence of all live cells. Data show mean ± standard error of three replicates. The dotted line indicates mean background fluorescence of untransfected cells.

### The DAD2 conformational biosensor retains catalytic activity

Ideally, the DAD2 conformational biosensors would retain SL binding and hydrolysis activities similar to the wild-type receptor, thus enabling a close to native condition direct-detection method (Figure 5A). To test enzymatic activity of DAD2cpGFP(G166), purified protein was incubated with *rac*-GR24, and both *rac*-GR24 and a hydrolysis product, ABC-FTL, were monitored by LC-MS/MS. The *rac*-GR24 peak intensity decreased in samples incubated with wild type DAD2 (DAD2wt) or DAD2cpGFP(G166) compared to a buffer-only control, and a new peak consistent with the fragmentation pattern of ABC-FTL^28^ appeared (Figure 5B). To confirm the identity of ABC-FTL, accurate mass was confirmed to within 1 ppm for both DAD2wt and DAD2cpGFP(G166) samples, and the fragmentation pattern matched that published previously ^28^ (Figure 5C).

**Figure 5.**
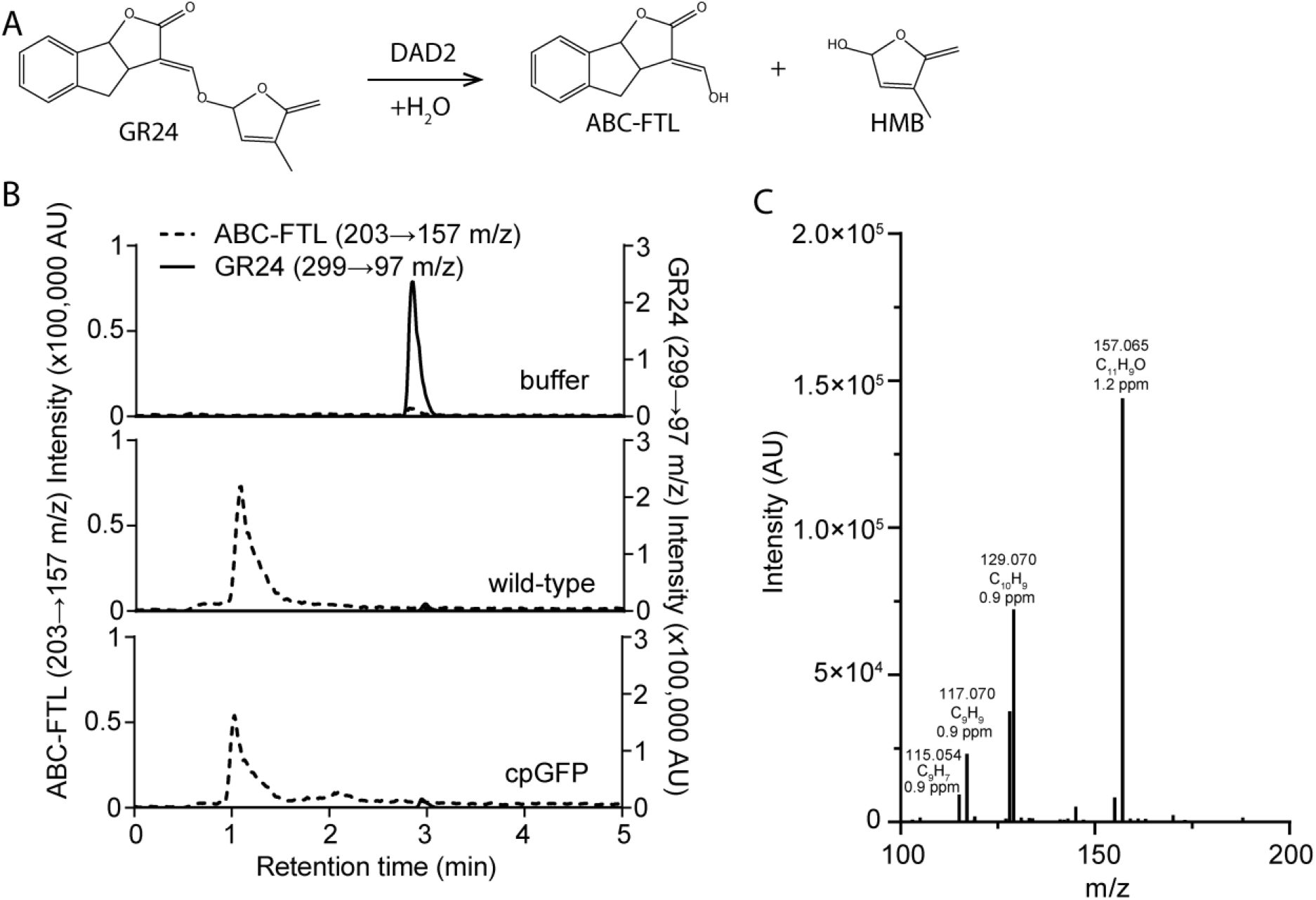
DAD2cpGFP(G166) hydrolysis activity towards *rac*-GR24. (A) DAD2 hydrolyses GR24 to produce two products, ABC-FTL and HMB. (B) Extracted ion chromatograms showing hydrolysis of *rac*-GR24 (299-›97 m/z transition, 3.0 min retention time) by wild-type DAD2 and DAD2cpGFP(G166) to form ABC-FTL (203-›157 m/z transition, 0.6 min retention time). Data show representative peak from three replicate samples. (C) MS2 Product ion scan of 203.07 m/z peak analysed on an Orbitap Elite mass spectrometer.

### Generation of an SL biosensor by FP domain insertion into ShHTL7

The different activities of SLs as hormones, parasitic seed germination stimulants, and mycorrhizae communication molecules are mediated by different receptors ^45^. In model plants, SLs are detected by D14/DAD2, whereas seed germination of the parasitic *S. hermonthica* is mediated by a related clade of α/β hydrolases that underwent convergent evolution to recognise SLs ^46^. The *S. hermonthica* HTL7 (ShHTL7) is the most sensitive of these receptors, and plays a dominant role in *S. hermonthica* seed germination ^47^. We asked whether we could apply the same principle used for DAD2cpGFP(G166) to generate an ShHTL7-based SL biosensor with the goal of being able to differentially detect the propensity of a SL or SL analogue to activate hormonal and parasitic seed germination pathways.

To rationally design an ShHTL7 version of the SL biosensor using DAD2cpGFP(G166) as a template, we aligned structures of ShHTL7 and DAD2 (PDB ID 5Z7Y ^48^ and 4DNP ^28^, respectively). The two receptors have conserved global tertiary structure (Figure 6A). We therefore hypothesised that domain insertion into the loop between helices 6 and 7 of ShHTL7 would facilitate coupling of SL-binding and cpGFP fluorescence, as observed in DAD2cpGFP(G166). This targeted approach allowed testing of fewer insertion variants: cpGFP was inserted into only four loop positions between helices 6 and 7. Again, LSSmOrange was fused to the C terminus to enable ratiometric measurement. Insertion after L166 displayed sensitivity to SL-binding, with greater than two-fold reduction in cpGFP fluorescence in the presence of 10 μM *rac*-GR24, and dose-responsive behaviour towards *rac*-GR24 (**Figure 6B, C**, Supplementary table 1). L166 is three amino acids C-terminal of the G166 insertion site in DAD2 based on sequence alignment, which highlights the importance of testing a library of insertion sites. Nevertheless, this again demonstrates the generalizable potential of domain insertion, and demonstrates that paralogous proteins can be successfully used as guides to narrow down library design.

**Figure 6.**
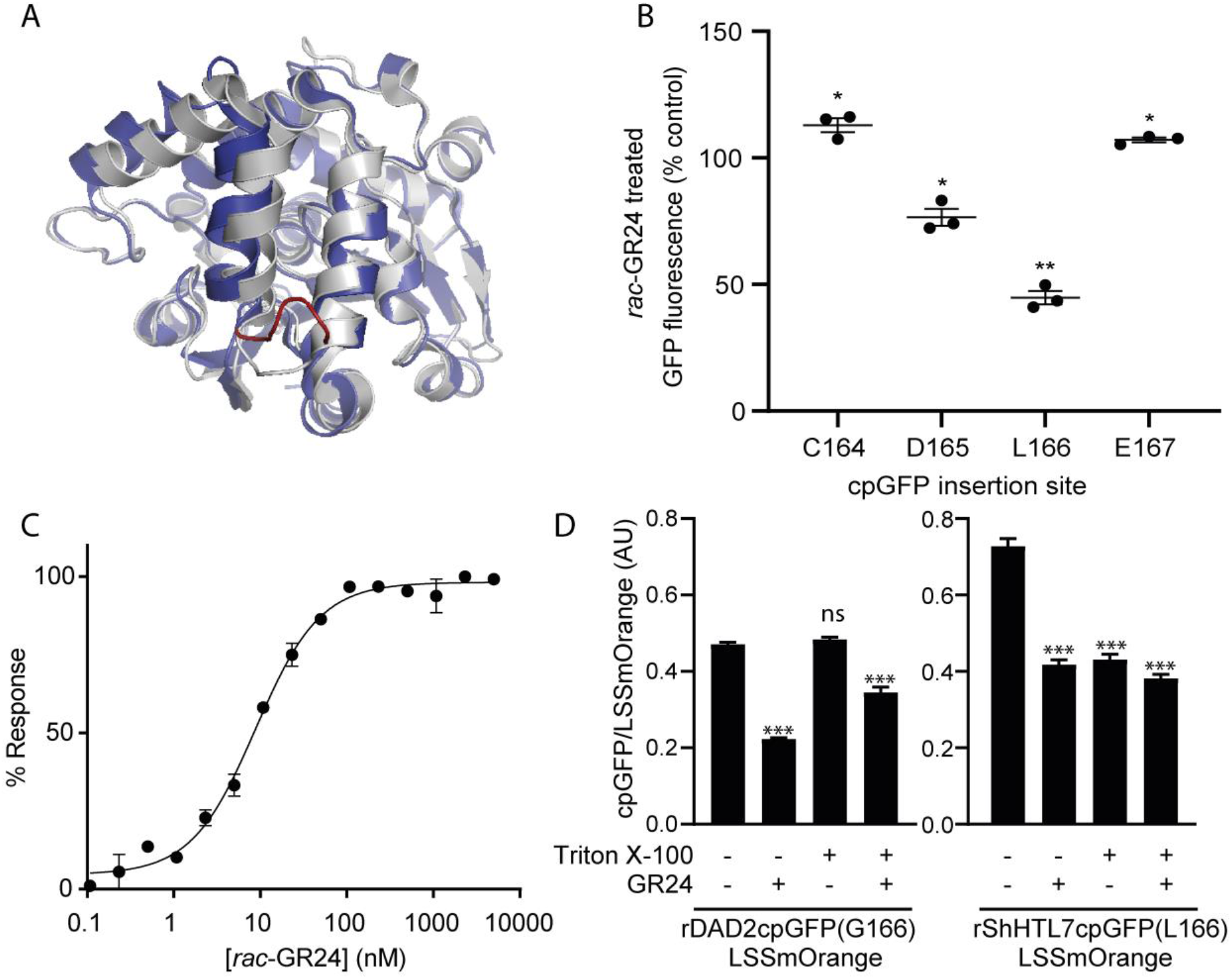
ShHTL7-based SL sensor using cpGFP loop insertion (A) Alignment of DAD2 (PDB ID 4DNP; grey) and ShHTL7 (PDB ID 5Z7Y; blue). cpGFP insertion sites in ShHTL7 are highlighted in red. (B) Fluorescent response to the strigolactone analogue *rac*-GR24 (10 μM) at each insertion site. ANOVA test for equality of means p<0.0001; *p<0.05, **p<0.01, one sample t-test compared to theoretical mean of 100 (no difference). C) Dose-response curve of rShHTL7cpGFP(L166) fused to *rac*-GR24 after 50 min incubation. (D) Fluorescent response of rDAD2cpGFP(G166) (left) and rShHTL7cpGFP(L166) (right) to 100 μM triton X-100 and 10 μM *rac*-GR24. Dunnett’s multiple comparisons test was used to compare each treatment condition to the untreated control: ns, not significant; ***p<0.001,. All error bars represent standard error of the mean of three replicates.

To assess whether rShHTL7cpGFP(L166) and rDAD2cpGFP(G166) retain the specificity of the parent receptors, we tested the effect of triton X-100 pre-treatment on each biosensor. Triton X-100 specifically inhibits ShHTL7 but not DAD2, due to a slightly more open position of helix α4 at the binding pocket entrance ^47^. When triton X-100 was pre-incubated with rDAD2cpGFP(G166) for two hours, it had no effect on the control lysate, and did not prevent *rac*-GR24-induced loss of cpGFP fluorescence (Figure 6D, Supplementary table 1). In contrast, triton X-100 reduced the cpGFP fluorescence of rShHTL7cpGFP(L166), and no further change was observed upon *rac*-GR24 treatment, consistent with triton X-100 binding and inhibition of ShHTL7cpGFP(L166)-LSSmOrange. Since ShHTL7 undergoes only minor conformational change upon triton X-100 binding, these results suggest that the cpGFP is sensitive to small local conformational changes.

## Discussion

Two new high-sensitivity biosensors were developed based on DAD2 and ShHTL7. The sensors provide an alternate and direct method of detection and quantitation for SLs, with low nanomolar sensitivity. The sensitivity is on par with current mass spectrometry limits of detection: a recent SL quantification method development paper reported a limit of detection of 4.94 μg/L (16.6 nM) for *rac*-GR24 ^49^. Furthermore, the sensors do not require specialised equipment, and are adaptable to high-throughput applications. Unlike mass spectrometry, the biosensors cannot identify specific strigolactones; however, given they report on receptor binding, they may be able to distinguish functionally active strigolactones from inactive precursors and other molecules. In this respect, the biosensors could be used as an alternative to existing functional assays, such as seed germination and branching inhibition bioassays, for some research applications. Seed germination assays in particular require alternatives as they pose environmental and agricultural risks associated with accidental release of parasitic seeds. As such, the biosensors are a valuable addition to the SL detection toolbox, complementing existing methods, with particular value in any high-throughput applications.

Biosensors have demonstrated utility in drug discovery screens due to their high specificity and affinity, and integrated output module. Similarly, the SL biosensors developed here could be used for screening small molecule libraries for agonists of DAD2 and/or ShHTL7. The specific binding of triton X-100 to ShHTL7 but not DAD2 sensors indicated that the selectivity of the two receptors was retained. Therefore, the sensors could be used to distinguish compounds that bind the hormonal SL receptor from those that bind the *S. hermonthica* seed germination receptor. An expected challenge in using SLs for agricultural applications is the risk of unintended activities arising from the diverse functions of SLs in hormonal, symbiotic and parasitic signalling. Increasing the specificity of the compounds towards one receptor type would reduce off-target effects. For example, field treatment with SL mimics to trigger suicidal seed germination is under investigation as a control method for *Striga* and *Orobanche spp.*. Compounds that trigger parasite seed germination while lacking hormonal activity would be preferred to avoid altering crop development. Conversely, agonists of DAD2 but not ShHTL7 would be preferred as plant protection products (for protection from drought stress for example), as they would not trigger potentially destructive parasitic seed germination. Compounds with selective hormonal or seed germination activity are already being identified, and the application of these biosensors may expedite similar studies ^50,51^.

The DAD2 conformational biosensor may also be useful to complement existing biosensors for studying SL signalling, and in particular for dissecting aspects of SL signal transduction. Existing SL biosensors report on degradation of D14 or D53, and as such, are dependent on many cellular processes, including the activity of additional signalling components, proteasomal degradation, and in the case of the D14-based biosensor, potentially on changes in gene transcription. As the only direct-detection biosensor developed to-date, the DAD2 conformational biosensor provides new potential for detecting SLs in plants and unravelling the processes that contribute to SL signalling. Demonstration of activity in tobacco protoplasts provides proof-of-concept for application of this biosensor in signalling studies.

Considerable differences in sensitivity were seen across the strigolactone biosensors developed to-date. StrigoQuant, which reported on D53 degradation, had very high sensitivity, detecting 5-deoxystrigol at concentrations as low as 100 fM, and orobanchol down to 10 nM ^26^. In contrast, the D14 degradation-based *in planta* bioassay had an EC50 of 1.63 μM GR24 ^27^. The conformational biosensors developed here had intermediate sensitivities, with EC50s in the low- to mid-nanomolar range, while transcriptional Y2H-based biosensors had lower sensitivity. These differences may be expected due to the differing reliance of each biosensor on additional cellular processes. For example, in the StrigoQuant system, ubiquitination of multiple luciferase-tagged D53 molecules by a single SL-induced SCF E3 ligase complex may mediate signal amplification. Conversely, the uncharacterised signalling steps between SL binding and D14 degradation may dampen signal. In addition, differences in the sensitivity of the parent receptors used, and changes induced through modification for biosensor construction, likely impact SL binding affinity.

D14/DAD2 and ShHTL7 are non-canonical receptors in that they hydrolyse strigolactones, and therefore also act as enzymes ^40^. Receptors, and more broadly small molecule binding proteins such as calmodulin and periplasmic binding proteins, have been widely used as biosensor components ^31–34^. In contrast, enzymes have been largely overlooked as input domains. To our knowledge, where enzyme biosensors have been developed previously, they have been either conformational biosensors, or sensors of non-substrate ligands.

There are several potential reasons for which enzymes may have traditionally been overlooked as biosensor input domains. Firstly, enzymes can have comparatively small conformational change upon substrate binding compared to receptors. However, it is becoming clear that small conformational changes can induce signal output (a phenomenon supported by the triton X-100 data presented here). Secondly, the temporary nature of substrate binding during enzyme catalysis could generate a transient signal that would be difficult to detect. Yet we found that signal from the DAD2 and ShHTL7 enzyme biosensors persisted sufficiently to measure fluorescence change even at low substrate concentrations. Although D14/DAD2 was initially proposed to form a stable complex with one of its hydrolysis products, more recent data demonstrated release of both products at an equally fast rate. Therefore, it is unlikely that these enzymes are unique in their ability to be adapted as fluorescent biosensors.

This work demonstrates the utility of enzymes as biosensors for their substrate. Previously, development of biosensors for new analytes relied on scouring databases for potential binding proteins or mutating known binding proteins to change their ligand specificity. However, many potential analytes are substrates for known enzymes that could be harnessed for biosensor development, significantly broadening the pool of characterised proteins from which we can draw input domains. Application of enzymes as biosensor recognition modules for their substrates sets a new precedent in biosensor design.

As these are the first biosensor designs, there is much scope to improve signal dynamic range via linker engineering and optimisation of fluorescent proteins. Furthermore, development of DAD2 and ShHTL7 biosensors with spectrally resolved fluorescent proteins may allow both biosensors to be used in a single-pot assay.

## Methods

### Plasmid construction

DAD2, ShHTL7, cpGFP and LSSmOrange were codon optimised and synthesised as gene fragments, where DAD2 was synthesised in fusion to LexA DNA binding domain for use in the Y2H-based sensor (Integrated DNA Technologies; supplementary table 2). DAD2 cpGFP variants were constructed by amplifying variable N- and C-terminal portions of each receptor, and assembling with the cpGFP fragment (with or without LSSmOrange) into pET19b between BlpI and NcoI using Gibson assembly (NEB). All primers are listed in supplementary table 3, and all PCR amplifications used Phusion high-fidelity polymerase as per the manufacturer’s protocol. N-terminal portions were amplified using primer DAD2N-F1 (DAD2 construct) and a variable reverse primer, whereas C-terminal portions were amplified using a variable forward primer and DAD2C-R1 (single-fluorophore DAD2 constructs) or DAD2C-R2 (LSSmOrange fusion DAD2 constructs). The LSSmOrange coding sequence was amplified using LSSmOrange-F and LSSmOrange-R primers. For ShHTL7 constructs, a base plasmid was initially constructed by assembling the ShHTL7 gene fragment with the LSSmOrange amplicon (LSSmOrange-F and LSSmOrange-R primers) into pET19b between BlpI and NcoI sites. The entire base plasmid was then amplified to introduce homologous ends to cpGFP at each insertion location using different primer sets (supplementary table 3), and the amplicons were assembled with the cpGFP gene fragment using Gibson cloning (NEB). 6His-DAD2 and 6His-DAD2cpGFP(G166) were amplified from DAD2 gene fragment and pET19b-DAD2cpGFP(G166), respectively, using primers DAD2N-F2 (1/10000 dilution of standard primer concentration), DAD2N-F3 and DAD2C-R1, where DAD2N-F3 anneals to the 5′ end of DAD2N-F2. This allowed initial amplification of the template by DAD2N-F2, followed by subsequent amplification by DAD2N-F3. Amplicons were cloned by Gibson assembly (NEB) into pET19b between BlpI and NcoI sites. PCR reactions used Phusion polymerase (NEB) or PrimeStarGXL polymerase (Takara) as per the manufacturers’ protocols.

For protoplast experiments, rDAD2cpGFP(G166) was amplified using primers 441 and 442, then digested with ClaI and SpeI and ligated into the minimal protoplast vector developed by Pouvreau *et al* ^52^.

pY2H-GFP was constructed by homologous recombination *in vivo* assembly in CEN.PK2-1C. Homologous arms for recombination at the YARCΔ8 locus were PCR amplified from CEN.PK113-7D genomic DNA using primers 101+102 for the upstream arm, and primers 122+123 for the downstream arm. PGAL1 and PGAL2 promoters and TIDP1T, TLCS2 and TRPL41B terminators were PCR amplified from CEN.PK113-7D genomic DNA using the following primer pairs: 103+104, 33+108, 107+32, 111+112, and 120+121 respectively. The GFP coding sequence was amplified from plasmid pILGFP3 (Addgene 65451) using primers 117+118. Coding sequences of a fusion of DAD2 to the LexA DNA binding domain (LexA-DAD2), and of MAX2 to the B42 activation domain (B42AD-MAX2) were synthesised as gene fragments (Integrated DNA Technologies). A synthetic promoter (PLexA) consisting of eight repeats of the LexA operator and a minimal GAL1 promoter was also synthesised (GenScript). LexA-DAD2, B42AD-MAX2 and PLexA were PCR amplified using primers 105+106, 109+110, and 113+124, respectively. All PCR fragments were purified using EasyPure ® PCR Purification Kit (TransGen Biotech) as per the manufacturers protocol and co-transformed into CEN.PK2-1C with pRS425 episomal plasmid backbone digested with SfoI and SacI (NEB). The assembled plasmid was extracted from S. cerevisiae, propagated in E. coli and verified by sequencing. Next, pY2H-GFP was adapted to include a URA3 selection marker to produce pY2H-GFP(URA3). GFP-TRPL41B and the downstream YARCΔ8 homolgous recombination are were PCR amplified from pY2H-GFP with primers 183+184 and 187+188, respectively. The URA3 selection marker was PCR amplified from pRS426 with primers 185+186. The resulting PCR fragments were re-assembled into pY2H-GFP digested with BssHII and NdeI using Gibson cloning (NEB) and confirmed by sequencing. The verified plasmid was digested with SwaI and transformed into CEN.PK2-1C for genomic integration at the YARCΔ8 locus, producing yeast strain ScY2H-GFP(URA3).

### Domain insertion screen

*E. coli* Rosetta (DE3) (DAD2 constructs) or BL21 (DE3) containing pGro7 (Takara) (ShHTL7 constructs) were used for recombinant protein expression. Fresh LB medium (10 mL) containing 100 μg/mL ampicillin and 34 μg/mL chloramphenicol was inoculated with overnight pre-cultures and grown at 37 °C, 200 rpm until 0.5 < OD_600 nm_<1.0. Cultures were induced with 1 mM IPTG then incubated for 16-20 h at 20 °C, 200 rpm. 2 mL aliquots of culture were centrifuged and *E. coli* were stored as pellets at −30 °C until use. *E. coli* were lysed by sonication (3 × 15 sec at 30% power) in 600 μL 50 mM Tris-HCl, 150 mM NaCl (pH 7.5) supplemented with 1 mg/mL lysozyme, 1 mM DTT and 1 × EDTA-free protease inhibitor cocktail (Roche). Lysates were clarified by centrifugation at 17,000 ×g, 4 °C for 10 min, then 200 μL supernatant was transferred to a microtitre plate containing *rac*-GR24 or solvent only control to a final concentration of 0.1 % acetone ± 10 μM *rac*-GR24. Fluorescence was measured using a Tecan Infinite 200 Pro microplate reader at 465 nm excitation, 517 nm emission (GFP) and 465 nm excitation, 575 nm emission (LSSmOrange).

### Strigolactone and triton X-100 treatments

Lysates expressing rDAD2cpGFP(G166) or rShHTL7cpGFP(G166) were prepared as per the domain insertion screen. For dose-response curves, 90 μL supernatant was transferred to a microtitre plate, followed by 10 μL of 2.15 nM – 100 μM *rac*-GR24serial dilutions in 50 mM Tris-HCl, 150 mM NaCl, 1% acetone (pH 7.5) to final concentrations 215 pM – 10 μM *rac*-GR24, 0.1% acetone after addition to lysate. Fluorescence was measured immediately and after 1 hour using a Tecan Infinite 200 Pro microplate reader as per the domain insertion screen. For triton X-100 analyses, clarified lysates were incubated for 2 h on ice with or without 100 μM triton X-100, then treated with 10 μM *rac*-GR24 or acetone control and fluorescence was measured immediately.

### Yeast cell culture and *rac*-GR24 treatment of yeast biosensor

During strain construction and all pre-cultures, yeast strains were grown on YPD, or where auxotrophic selection was required, on 6.7 g/L yeast nitrogen base without amino acids (YNB) supplemented with 20 g/L glucose. YNB pH was adjusted to 6.0 with ammonium hydroxide, and YNB was supplemented with 20 mg/L histidine, 100 mg/L leucine and 100 mg/L tryptophan (ScY2H-GFP(URA3)), and with 20 mg/L uracil for the CEN.PK2-1C control. For cultivation on petri dishes, media were solidified with 20 g/L bacteriological agar.

For *rac*-GR24 treatment experiments, ScY2H-GFP(URA3) yeast were streaked from glycerol stocks onto YNB agar and grown for 2 days at 30 °C. Single colonies were inoculated into a primary pre-culture in 5 mL YNB in McCartney bottles and grown for 24 hours. A secondary pre-culture (5 mL YNB in a McCartney bottle) was inoculated from the primary pre-culture using a volume of culture such that the secondary pre-culture would reach mid-log phase the following morning (based on cell density and preliminary growth rate estimates). Experimental cultures were inoculated from secondary pre-cultures to a starting culture density of OD600nm=0.2 into 100 mL shake flasks containing 5 mL YNB MES ++ medium (6.7 g/L YNB without amino acids, 100 mM MES pH 6.0, 100 mg/L glutamic acid, 120 mg/L lysine, 40 mg/L methionine, 50 mg/L phenylalanine, 375 mg/L serine, 200 mg/L threonine, and 10 mg/L myo-inositol) supplemented with 2% galactose, 35 mg/L histidine, 110 mg/L leucine, 100 mg/L tryptophan, and 40 mg/L uracil (CEN.PK2-1C only), with or without 10 μM *rac*-GR24. Strains were cultured for 16 h at 30 °C, 200 rpm, then GFP fluorescence was measured by flow cytometry (BD Accuri™ C6; BD Biosciences, USA) using a 488 nm laser and 530/20 nm band-pass filter. The dose-response curves were performed as described with the exception that experimental cultures were in 1 mL medium, in 6-well plates.

### Protein purification

6His-DAD2 and 6His-DAD2cpGFP(G166) were expressed as per the domain insertion screen, except 800 mL cultures were used, and expression was induced with 400 μM IPTG. *E. coli* were lysed in 10 mM Hepes-NaOH, 150 mM NaCl (pH 7.5) by passing twice trough an Emulsiflex C5 high pressure homogeniser (Avestin) at 15,000 psi, then clarified by centrifugation at 18,000 ×g 4 °C for 30 min. Supernatants were filtered through 0.22 μm polyethersulfone Millex-GP syringe filters, then supplemented to a final concentration 20 mM imidazole. Each protein was purified using a 1 mL HisTrap HP column (GE Healthcare), washing with 10 mM Hepes-NaOH, 150 mM NaCl, 20 mM imidazole (pH7.5) and eluting with 10 mM Hepes-NaOH, 150 mM NaCl, 200 mM imidazole (pH7.5). Protein concentration was quantified using a NanoDrop.

### GR24 hydrolysis assays

30 μM 6His-DAD2, 6His-DAD2cpGFP(G166) or a buffer-only control was incubated with 30 μM *rac*-GR24 in 10 mM Hepes-NaOH, 150 mM NaCl pH 7.5 in 50 μL reaction volumes at 30 °C for 2 h. *rac*-GR24 and reaction products were extracted twice into 50 μL ethyl acetate. The extract was diluted with 100 μL acetonitrile then vacuum centrifuged to 30 μL. 2 μL of each extraction was analysed by LC-MS/MS. The LC-MS system was a Nexera X2 ultra high pressure liquid chromatograph (UHPLC) system (Shimadzu Corporation, Kyoto, Japan) coupled with a 5500 QTRAP MS system equipped with an electrospray ionization source (ESI) (AB Sciex, USA). SLs were separated on a Phenomenex Kinetex C18 reversed phase column (2.1mm × 100 mm, 1.7 μm) maintained at 45 °C. The settings and gradients for UPLC are as follows: flow rate: 0.5 mL/min; mobile phase A:0.1 % acetic acid in deionized water; mobile phase B: 0.1 % acetic acid in acetonitrile; The gradient was: 30 % to 100 % B for 12 minutes, followed by 3 minutes wash with 100 % B. MS/MS spectra were recorded using multiple reaction monitoring with the following settings: scan mode: positive; curtain gas: 20 psi; collision gas: high; ion spray voltage: 5500 V; ion source temperature: 500 °C; ion source gas 1: 25 psi; ion source gas 2: 25 psi. Declustering potential, entrance potential, collision energy and collision cell exit potential were set to 80 V, 10 V, 20 V and 15 V, respectively.

Accurate mass and fragmentation patterns were confirmed as follows. Samples were separated using reversed-phase chromatography on a Dionex Ultimate 3000 RSLC nano-system. Using a flow rate of 30 μl/min, samples were desalted on a Thermo PepMap 100 C18 trap (0.3 × 5 mm, 5 μm) for 5 min, followed by separation on a Vydac Everest C18 column (150 mm × 75 μm, 5um particle) at a flow rate of 300 nL/min. A gradient of 10-50% buffer B over 7 min where buffer A + 1 % ACN / 0.1% FA and buffer B + 80% ACN / 0.1% FA was used to separate molecules, followed by washing at 98% B for 3 min, before returning to 10% B . Eluted molecules were directly analysed on an Orbitap Elite mass spectrometer (Thermo) using an NSI electrospray interface. Source parameters included a capillary temperature of 275° C; S-Lens RF level at 60%; source voltage of 2.4 kV and maximum injection times of 200 ms for MS and 150 ms for MS2. Instrument parameters included an FTMS scan across m/z range 50-500 at 60,000 resolution followed by information dependent acquisition of the top 10 singly charged peaks in the Orbitrap using HCD fragmentation, and normalised collision energy of 64 V. Dynamic ion exclusion was employed using a 15-second interval. Charge state screening was enabled with rejection of all ions other than +1 charged species.

### Protoplast preparation and analysis

Tobacco protoplasts were prepared as per Pouvreau et al ^52^. Flow cytometry analyses were conducted on an Invitrogen Attune NxT Flow Cytometer and analysed with Invitrogen Attune NxT Software. Cells suspended in the protoplast buffer at a concentration of 0.187 M cells per ml were analysed at a flow rate of 200 μl/min and data were acquired for 150 μl. Sample were complemented with propidium iodide (PI) at 1 μg/ml final concentration and incubated 15 min prior to Flow cytometric analyses. Chlorophyll and PI fluorescence were excited with a 561-nm laser and emissions were respectively captured by a 780/60- nm and a 620/ 15-nm band-pass filters. Protoplast population was gated based on SSC/FSC as well as Chlorophyll fluorescence/FSC ratios based on previous data (data not shown). Positive cells for PI staining (regarded as dead cells) were ignored for the rest of the analyses. For the remaining cells, LSSmOrange and cpGFP fluorescence were excited with a 488-nm laser and emissions were respectively captured by a 590/ 40-nm and a 530/ 30-nm band-pass filters. Unless otherwise stated, analyses were conducted on a minimum of three individual samples for each condition.

Each sample was run once through the flow cytometer and then immediately spiked 1:100 with GR4 in 10 % acetone solution. Eight different final GR4 dilutions were used: 10 uM, 1 uM, 100 nM, 31.6 nM, 10 nM, 1 nM, 0.1 nM, and 0 nM. Samples were then run twice more through the flow cytometer at 30 minutes interval.

### Statistical analyses

All statistical analyses were performed in GraphPad Prism 8. For insertion site screening, the differences in cpGFP fluorescence between GR24 and control were tested using one sample t-tests (two-tailed). To test responses to Triton X-100, data were compared using one-way analysis of variance (ANOVA). Post-hoc Dunnett’s multiple comparison tests were used to compare each treatment against the –Triton X-100 –GR24 control. For *in vitro* dose-response curves, GFP/LssmOrange fluorescence intensities were normalised to a scale of 0 to 100, and fit to an [Agonist] vs. response – Variable slope (four parameters) model. Where EC50 values were compared, an extra sum-of-squares F test was used, with a P value cutoff of 0.05. *In vivo* protoplast or yeast dose-response curves were fit to an [Agonist] vs. response – Variable slope (four parameters) model.

## Acknowledgements

The authors acknowledge the facilities, and the scientific and technical assistance of the School of Chemistry and Molecular Biosciences Mass spectrometry facility and The University of Queensland. R.J.C is funded by a CSIRO Synthetic Biology Future Science Platform PhD Top-Up Scholarship, and an Australian Government Research Training Program Stipend. J.H.W. is supported by a CSIRO Synthetic Biology Future Fellowship. B.P. was partly funded through the CSIRO Synthetic Biology Future Science Platform and the CSIRO Research Office CERC Postdoctoral Fellowship scheme. C.A.B. is funded by an ARC Georgina Sweet Laureate Fellowship FL180100139.

## Author contributions

R.J.C. designed and performed experiments and interpreted data. J.H.W. developed the biosensor designs and oversaw the study design. R.J.C. and J.H.W. wrote the manuscript. B.P. performed all protoplast experiments. D.C. developed methods for analysis of GR24 hydrolysis and did critical revision of the manuscript. C.A.B. and C.E.V. critical reviewed the manuscript and advised on study design.

## References

(1) Agusti, J. Strigolactone-Mediated Stimulation of Secondary Xylem Proliferation in Stems. Xylem Methods Protoc. 2017, 21–26.

(2) Gomez-Roldan, V.; Fermas, S.; Brewer, P. B.; Puech-Pagès, V.; Dun, E. A.; Pillot, J.-P.; Letisse, F.; Matusova, R.; Danoun, S.; Portais, J.-C. Strigolactone Inhibition of Shoot Branching. Nature 2008, 455 (7210), 189–194.

(3) Kapulnik, Y.; Delaux, P.-M.; Resnick, N.; Mayzlish-Gati, E.; Wininger, S.; Bhattacharya, C.; Séjalon-Delmas, N.; Combier, J.-P.; Bécard, G.; Belausov, E. Strigolactones Affect Lateral Root Formation and Root-Hair Elongation in Arabidopsis. Planta 2011, 233 (1), 209–216.

(4) Rasmussen, A.; Mason, M. G.; De Cuyper, C.; Brewer, P. B.; Herold, S.; Agusti, J.; Geelen, D.; Greb, T.; Goormachtig, S.; Beeckman, T. Strigolactones Suppress Adventitious Rooting in Arabidopsis and Pea. Plant Physiol. 2012, 158 (4), 1976–1987.

(5) Umehara, M.; Hanada, A.; Yoshida, S.; Akiyama, K.; Arite, T.; Takeda-Kamiya, N.; Magome, H.; Kamiya, Y.; Shirasu, K.; Yoneyama, K. Inhibition of Shoot Branching by New Terpenoid Plant Hormones. Nature 2008, 455 (7210), 195–200.

(6) Yamada, Y.; Furusawa, S.; Nagasaka, S.; Shimomura, K.; Yamaguchi, S.; Umehara, M. Strigolactone Signaling Regulates Rice Leaf Senescence in Response to a Phosphate Deficiency. Planta 2014, 240 (2), 399–408.

(7) Davidson, E. A.; Bayer, T. S.; Windram, O.; Hleba, Y. Strigolactone Formulations and Uses Thereof. US20160159780, 2016.

(8) Van Ha, C.; Leyva-González, M. A.; Osakabe, Y.; Tran, U. T.; Nishiyama, R.; Watanabe, Y.; Tanaka, M.; Seki, M.; Yamaguchi, S.; Van Dong, N. Positive Regulatory Role of Strigolactone in Plant Responses to Drought and Salt Stress. Proc. Natl. Acad. Sci. 2014, 111 (2), 851–856.

(9) Liu, J.; He, H.; Vitali, M.; Visentin, I.; Charnikhova, T.; Haider, I.; Schubert, A.; Ruyter-Spira, C.; Bouwmeester, H. J.; Lovisolo, C. Osmotic Stress Represses Strigolactone Biosynthesis in Lotus Japonicus Roots: Exploring the Interaction between Strigolactones and ABA under Abiotic Stress. Planta 2015, 241 (6), 1435–1451.

(10) Akiyama, K.; Matsuzaki, K.; Hayashi, H. Plant Sesquiterpenes Induce Hyphal Branching in Arbuscular Mycorrhizal Fungi. Nature 2005, 435 (7043), 824–827. https://doi.org/10.1038/nature03608.

(11) Vurro, M.; Boari, A.; Thiombiano, B.; Bouwmeester, H. Strigolactones and Parasitic Plants. In Strigolactones - Biology and Applications; Springer International Publishing: Cham, 2019; pp 89–120. https://doi.org/10.1007/978-3-030-12153-2_3.

(12) Parker, C. Observations on the Current Status of Orobanche and Striga Problems Worldwide. Pest Manag. Sci. 2009, 65 (5), 453–459.

(13) Kgosi, R. L.; Zwanenburg, B.; Mwakaboko, A. S.; Murdoch, A. J. Strigolactone Analogues Induce Suicidal Seed Germination of Striga Spp. in Soil. Weed Res. 2012, 52 (3), 197–203.

(14) Samejima, H.; Babiker, A. G.; Takikawa, H.; Sasaki, M.; Sugimoto, Y. Practicality of the Suicidal Germination Approach for Controlling Striga Hermonthica. Pest Manag. Sci. 2016.

(15) Zwanenburg, B.; Mwakaboko, A. S.; Kannan, C. Suicidal Germination for Parasitic Weed Control. Pest Manag. Sci. 2016.

(16) Vurro, M.; Prandi, C.; Baroccio, F. Strigolactones: How Far Is Their Commercial Use for Agricultural Purposes? Pest Manag. Sci. 2016.

(17) Gobena, D.; Shimels, M.; Rich, P. J.; Ruyter-Spira, C.; Bouwmeester, H.; Kanuganti, S.; Mengiste, T.; Ejeta, G. Mutation in Sorghum LOW GERMINATION STIMULANT 1 Alters Strigolactones and Causes Striga Resistance. Proc. Natl. Acad. Sci. 2017, 114 (17), 4471–4476.

(18) Dor, E.; Yoneyama, K.; Wininger, S.; Kapulnik, Y.; Yoneyama, K.; Koltai, H.; Xie, X.; Hershenhorn, J. Strigolactone Deficiency Confers Resistance in Tomato Line SL-ORT1 to the Parasitic Weeds Phelipanche and Orobanche Spp. Phytopathology 2011, 101 (2), 213–222. https://doi.org/10.1094/PHYTO-07-10-0184.

(19) Fernández-Aparicio, M.; Kisugi, T.; Xie, X.; Rubiales, D.; Yoneyama, K. Low Strigolactone Root Exudation: A Novel Mechanism of Broomrape (Orobanche and Phelipanche Spp.) Resistance Available for Faba Bean Breeding. J. Agric. Food Chem. 2014, 62 (29), 7063–7071. https://doi.org/10.1021/jf5027235.

(20) Pavan, S.; Schiavulli, A.; Marcotrigiano, A. R.; Bardaro, N.; Bracuto, V.; Ricciardi, F.; Charnikhova, T.; Lotti, C.; Bouwmeester, H.; Ricciardi, L. Characterization of Low-Strigolactone Germplasm in Pea (Pisum Sativum L.) Resistant to Crenate Broomrape (Orobanche Crenata Forsk.). Mol. Plant. Microbe. Interact. 2016, 29 (10), 743–749. https://doi.org/10.1094/MPMI-07-16-0134-R.

(21) Yoneyama, K.; Xie, X.; Yoneyama, K.; Kisugi, T.; Nomura, T.; Nakatani, Y.; Akiyama, K.; McErlean, C. S. P. Which Are the Major Players, Canonical or Non-Canonical Strigolactones? J. Exp. Bot. 2018, 69 (9), 2231–2239. https://doi.org/10.1093/jxb/ery090.

(22) Jia, K.-P.; Li, C.; Bouwmeester, H. J.; Al-Babili, S. Strigolactone Biosynthesis and Signal Transduction. In Strigolactones - Biology and Applications; Springer International Publishing: Cham, 2019; pp 1–45. https://doi.org/10.1007/978-3-030-12153-2_1.

(23) Seto, Y.; Yasui, R.; Kameoka, H.; Tamiru, M.; Cao, M.; Terauchi, R.; Sakurada, A.; Hirano, R.; Kisugi, T.; Hanada, A.; et al. Strigolactone Perception and Deactivation by a Hydrolase Receptor DWARF14. Nat. Commun. 2019, 10 (1), 191. https://doi.org/10.1038/s41467-018-08124-7.

(24) Chevalier, F.; Nieminen, K.; Sánchez-Ferrero, J. C.; Rodríguez, M. L.; Chagoyen, M.; Hardtke, C. S.; Cubas, P. Strigolactone Promotes Degradation of DWARF14, an α/β Hydrolase Essential for Strigolactone Signaling in Arabidopsis. Plant Cell 2014, 26 (3), 1134–1150. https://doi.org/10.1105/tpc.114.122903.

(25) Hu, Q.; He, Y.; Wang, L.; Liu, S.; Meng, X.; Liu, G.; Jing, Y.; Chen, M.; Song, X.; Jiang, L.; et al. DWARF14, a Receptor Covalently Linked with the Active Form of Strigolactones, Undergoes Strigolactone-Dependent Degradation in Rice. Front. Plant Sci. 2017, 8. https://doi.org/10.3389/fpls.2017.01935.

(26) Samodelov, S. L.; Beyer, H. M.; Guo, X.; Augustin, M.; Jia, K.-P.; Baz, L.; Ebenhöh, O.; Beyer, P.; Weber, W.; Al-Babili, S. StrigoQuant: A Genetically Encoded Biosensor for Quantifying Strigolactone Activity and Specificity. Sci. Adv. 2016, 2 (11), e1601266.

(27) Sanchez, E.; Artuso, E.; Lombardi, C.; Visentin, I.; Lace, B.; Saeed, W.; Lolli, M. L.; Kobauri, P.; Ali, Z.; Spyrakis, F.; et al. Structure-Activity Relationships of Strigolactones via a Novel, Quantitative in Planta Bioassay. J. Exp. Bot. 2018, 69 (9), 2333–2343. https://doi.org/10.1093/jxb/ery092.

(28) Hamiaux, C.; Drummond, R. S. M. S. M.; Janssen, B. J. J.; Ledger, S. E. E.; Cooney, J. M. M.; Newcomb, R. D. D.; Snowden, K. C. C. DAD2 Is an α/β Hydrolase Likely to Be Involved in the Perception of the Plant Branching Hormone, Strigolactone. Curr. Biol. 2012, 22 (21), 2032–2036. https://doi.org/10.1016/J.CUB.2012.08.007.

(29) Toh, S.; Holbrook-Smith, D.; Stokes, M. E. E.; Tsuchiya, Y.; McCourt, P. Detection of Parasitic Plant Suicide Germination Compounds Using a High-Throughput Arabidopsis HTL/KAI2 Strigolactone Perception System. Chem. Biol. 2014, 21 (8), 988–998. https://doi.org/10.1016/J.CHEMBIOL.2014.07.005.

(30) Tamura, T.; Hamachi, I. Recent Progress in Design of Protein-Based Fluorescent Biosensors and Their Cellular Applications. ACS Chem. Biol. 2014, 9 (12), 2708–2717. https://doi.org/10.1021/cb500661v.

(31) Nadler, D. C.; Morgan, S.-A.; Flamholz, A.; Kortright, K. E.; Savage, D. F. Rapid Construction of Metabolite Biosensors Using Domain-Insertion Profiling. Nat. Commun. 2016, 7 (1), 12266. https://doi.org/10.1038/ncomms12266.

(32) Ostermeier, M. Engineering Allosteric Protein Switches by Domain Insertion. Protein Eng. Des. Sel. 2005, 18 (8), 359–364. https://doi.org/10.1093/protein/gzi048.

(33) Marvin, J. S.; Schreiter, E. R.; Echevarría, I. M.; Looger, L. L. A Genetically Encoded, High-Signal-to-Noise Maltose Sensor. Proteins Struct. Funct. Bioinforma. 2011, 79 (11), 3025–3036. https://doi.org/10.1002/prot.23118.

(34) Hu, H.; Gu, Y.; Xu, L.; Zou, Y.; Wang, A.; Tao, R.; Chen, X.; Zhao, Y.; Yang, Y. A Genetically Encoded Toolkit for Tracking Live-Cell Histidine Dynamics in Space and Time. Sci. Rep. 2017, 7 43479. https://doi.org/10.1038/srep43479.

(35) Baird, G. S.; Zacharias, D. A.; Tsien, R. Y. Circular Permutation and Receptor Insertion within Green Fluorescent Proteins. Proc. Natl. Acad. Sci. 1999, 96 (20), 11241–11246. https://doi.org/10.1073/pnas.96.20.11241.

(36) Zhao, L.-H.; Zhou, X. E.; Wu, Z.-S.; Yi, W.; Xu, Y. Y.; Li, S.; Xu, T.-H.; Liu, Y.; Chen, R.-Z.; Kovach, A.; et al. Crystal Structures of Two Phytohormone Signal-Transducing α/β Hydrolases: Karrikin-Signaling KAI2 and Strigolactone-Signaling DWARF14. Cell Res. 2013, 23 (3), 436–439. https://doi.org/10.1038/cr.2013.19.

(37) Yao, R.; Ming, Z.; Yan, L.; Li, S.; Wang, F.; Ma, S.; Yu, C.; Yang, M.; Chen, L.; Chen, L.; et al. DWARF14 Is a Non-Canonical Hormone Receptor for Strigolactone. Nature 2016, 536. https://doi.org/10.1038/nature19073.

(38) Ast, C.; Foret, J.; Oltrogge, L. M.; De Michele, R.; Kleist, T. J.; Ho, C.-H.; Frommer, W. B. Ratiometric Matryoshka Biosensors from a Nested Cassette of Green- and Orange-Emitting Fluorescent Proteins. Nat. Commun. 2017, 8 (1), 431. https://doi.org/10.1038/s41467-017-00400-2.

(39) Bürger, M.; Chory, J. The Many Models of Strigolactone Signaling. Trends Plant Sci. 2020. https://doi.org/10.1016/j.tplants.2019.12.009.

(40) Yao, R.; Wang, F.; Ming, Z.; Du, X.; Chen, L.; Wang, Y.; Zhang, W.; Deng, H.; Xie, D. ShHTL7 Is a Non-Canonical Receptor for Strigolactones in Root Parasitic Weeds. Cell Res. 2017, 27 (6), 838–841. https://doi.org/10.1038/cr.2017.3.

(41) de Saint Germain, A.; Clavé, G.; Badet-Denisot, M.-A.; Pillot, J.-P.; Cornu, D.; Le Caer, J.-P.; Burger, M.; Pelissier, F.; Retailleau, P.; Turnbull, C.; et al. An Histidine Covalent Receptor and Butenolide Complex Mediates Strigolactone Perception. Nat. Chem. Biol. 2016, 12 (10), 787–794. https://doi.org/10.1038/nchembio.2147.

(42) Wang, L.; Wang, B.; Jiang, L.; Liu, X.; Li, X.; Lu, Z.; Meng, X.; Wang, Y.; Smith, S. M.; Li, J. Strigolactone Signaling in Arabidopsis Regulates Shoot Development by Targeting D53-like SMXL Repressor Proteins for Ubiquitination and Degradation. Plant Cell 2015, 27 (11), 3128–3142.

(43) Zhao, J.; Wang, T.; Wang, M.; Liu, Y.; Yuan, S.; Gao, Y.; Yin, L.; Sun, W.; Peng, L.; Zhang, W.; et al. DWARF3 Participates in an SCF Complex and Associates with DWARF14 to Suppress Rice Shoot Branching. Plant Cell Physiol. 2014, 55 (6), 1096–1109. https://doi.org/10.1093/pcp/pcu045.

(44) Song, X.; Lu, Z.; Yu, H.; Shao, G.; Xiong, J.; Meng, X.; Jing, Y.; Liu, G.; Xiong, G.; Duan, J.; et al. IPA1 Functions as a Downstream Transcription Factor Repressed by D53 in Strigolactone Signaling in Rice. Cell Res. 2017, 27 (9), 1128–1141. https://doi.org/10.1038/cr.2017.102.

(45) Lumba, S.; Holbrook-Smith, D.; McCourt, P. The Perception of Strigolactones in Vascular Plants. Nat. Chem. Biol. 2017, 13 (6), 599–606. https://doi.org/10.1038/nchembio.2340.

(46) Conn, C. E.; Bythell-Douglas, R.; Neumann, D.; Yoshida, S.; Whittington, B.; Westwood, J. H.; Shirasu, K.; Bond, C. S.; Dyer, K. A.; Nelson, D. C. Convergent Evolution of Strigolactone Perception Enabled Host Detection in Parasitic Plants. Science 2015, 349 (6247), 540–543. https://doi.org/10.1126/science.aab1140.

(47) Shahul Hameed, U.; Haider, I.; Jamil, M.; Kountche, B. A.; Guo, X.; Zarban, R. A.; Kim, D.; Al-Babili, S.; Arold, S. T. Structural Basis for Specific Inhibition of the Highly Sensitive ShHTL7 Receptor. EMBO Rep. 2018, 19 (9), e45619. https://doi.org/10.15252/embr.201745619.

(48) Xu, Y.; Miyakawa, T.; Nosaki, S.; Nakamura, A.; Lyu, Y.; Nakamura, H.; Ohto, U.; Ishida, H.; Shimizu, T.; Asami, T.; et al. Structural Analysis of HTL and D14 Proteins Reveals the Basis for Ligand Selectivity in Striga. Nat. Commun. 2018, 9 (1), 3947. https://doi.org/10.1038/s41467-018-06452-2.

(49) Boutet-Mercey, S.; Perreau, F.; Roux, A.; Clavé, G.; Pillot, J.-P.; Schmitz-Afonso, I.; Touboul, D.; Mouille, G.; Rameau, C.; Boyer, F.-D. Validated Method for Strigolactone Quantification by Ultra High-Performance Liquid Chromatography-Electrospray Ionisation Tandem Mass Spectrometry Using Novel Deuterium Labelled Standards. 2017. https://doi.org/10.1002/pca.2714.

(50) Uraguchi, D.; Kuwata, K.; Hijikata, Y.; Yamaguchi, R.; Imaizumi, H.; Am, S.; Rakers, C.; Mori, N.; Akiyama, K.; Irle, S.; et al. A Femtomolar-Range Suicide Germination Stimulant for the Parasitic Plant *Striga Hermonthica*. Science (80-.). 2018, 362 (6420), 1301–1305. https://doi.org/10.1126/science.aau5445.

(51) Yasui, R.; Seto, Y.; Ito, S.; Kawada, K.; Itto-Nakama, K.; Mashiguchi, K.; Yamaguchi, S. Chemical Screening of Novel Strigolactone Agonists That Specifically Interact with DWARF14 Protein. Bioorganic Med. Chem. Lett. 2019, 29 (7), 938–942. https://doi.org/10.1016/j.bmcl.2019.01.010.

(52) Pouvreau, B.; Blundell, C.; Vohra, H.; Zwart, A. B.; Arndell, T.; Singh, S.; Vanhercke, T. A Versatile High Throughput Screening Platform for Plant Metabolic Engineering Highlights the Major Role of ABI3 in Lipid Metabolism Regulation. bioRxiv 2019, 853606. https://doi.org/10.1101/853606.

