## Supplementary data for "Rational design of novel fluorescent enzyme biosensors for direct detection of strigolactones"

Supplementary table 1. Statistical analyses

**Insertion sites for cpGFP into DAD2 and ShHTL7**

One sample t-tests

| | A151 | G166 | A167 | N182 | M183 | R184 | C164 | D165 | L166 | E167 |
|---|---|---|---|---|---|---|---|---|---|---|
| t | 2.134 | 61.87 | 12.24 | 2.061 | 4.130 | 9.319 | 4.688 | 6.997 | 21.11 | 7.398 |
| df | 2 | 2 | 2 | 2 | 2 | 2 | 2 | 2 | 2 | 2 |
| p | 0.1665 | 0.0003 | 0.0066 | 0.1754 | 0.0539 | 0.0113 | 0.0426 | 0.0198 | 0.0022 | 0.0178 |

**Responses of rDAD2cpGFP(G166) and rShHTL7cpGFP(L166) to Triton X-100 and GR24**

One-way ANOVA

| | F | DFn | DFd | P |
|---|---|---|---|---|
| rDAD2cpGFP(G166) | 227.3 | 3 | 8 | <0.0001 |
| rShHTL7cpGFP(L166) | 117.5 | 3 | 8 | <0.0001 |

Dunnett's multiple comparisons tests

| rDAD2cpGFP(G166) | q | DF | p |
|---|---|---|---|
| -GR24-Triton vs. +GR24-Triton | 21.63 | 8 | <0.0001 |
| -GR24-Triton vs. -GR24+Triton | 1.089 | 8 | 0.5884 |
| -GR24-Triton vs. +GR24+Triton | 10.96 | 8 | <0.0001 |
| **rShHTL7cpGFP(L166)** | **q** | **DF** | **p** |
| -GR24-Triton vs. +GR24-Triton | 14.83 | 8 | <0.0001 |
| -GR24-Triton vs. -GR24+Triton | 14.21 | 8 | <0.0001 |
| -GR24-Triton vs. +GR24+Triton | 16.56 | 8 | <0.0001 |

Supplementary table 2: Coding sequences for DAD2, ShHTL7, cpEGFP and LSSmOrange

**LexA-DAD2**

atgaaagctctgacggctagacaacaggaggtgttcgatttgatcagagaccacatctctcaaacgggtatgccgccaacgagggccgaaatcgcccaaaga
ttaggattccgttctccaaatgcggccgaggagcatctaaaagcattagccagaaaaggtgtgatcgaaattgtgagcggagccagcagaggtattagattgct
tcaggaggaggaagaggggcttccattagttggcagggtagctgcaggcgagcctttgttagcccagcaacacatagagggccactaccaggtagacccct
cactgttcaagcccaacgcagatttcttgttgagagtcagtggcatgtccatgaaggatatcggtatcatggatggtgacttacttgccgtacacaagacgcaag
atgtacgtaacggccaggtggttgtggctcgtattgatgatgaagttactgtgaagaggttaaaaaaacagggcaacaaagtagaactacttcctgaaaattctg
agttcaagccaatagtggttgacttgagacaacagtcttttaccatagagggtttagctgttggagttatacgtaacggagactggctacctgccgcgaagaggg
tcaagctggatggcggaggtggttccggaggtggtggttccatgggtcaaactttactggacgccttgaatgtgagggttgtaggcagcggagagagagtgtt
ggtacttgctcacgggtttggtaccgaccagtccgcttggaataggattctacccttttccttagggactatagagttgtactatacgatttggtgtgtgcaggatca
gttaaccctgacttctttgattttaggaggtacacaacacttgacccttatgtggatgatttgctgcatatattagacgcgctggggatagactgttgcgcttatgttg
gtcattcagtatcagcaatgatcggcatattggcgtctatacgtaggccggaattattttctaagctaattttgataggcgcgagtcctaggtttctaaacgatgaag
actaccacggcggtttcgagcaaggagaaatcgaaaaggtcttttccgcgatggaggcaaactatgaggcgtgggtgaatggattcgcgcctttggcagtgg
gcgctgatgtaccggcagcggtaagggaattctcccgtacactgtttaacatgcgtccagacataactttgttcgttagtaggacagtattcaacagcgatatgag
aggtgttttgggcttagtgaaggtcccttgtcatatctttcagacagctagagatcattctgtgcctgcgagtgtagccacttatctaaagaaccacctaggcggta
aaaatacagtgcactggcttaacatagaaggtcaccttccacatctaagtgcgcccactcttttagcacaagagcttaggcgtgcactgtctcatcgttaa

**ShHTL7**

atgagtagcattgggttagcgcacaatgttacgatcctgggaagcggagaaactaccgtggtattgggtcacggatacggcactgatcagtctgtgtggaaact
gcttgtaccgtacttggttgatgattacaaggtcttactttacgatcatatgggggcgggcacaaccaaccctgattatttcgatttcgaccgctatagttcactgga
gggctacagttacgacttgatcgcaattttggaagagttccaagtatcgaagtgcatttacgttggacactcgatgtcttcgatggcagcggcagtcgcatctatttt
tcgccctgatctgtttcacaaattagtaatgatcagccccacaccccgccttattaacactgaagaatactacggtggatttgagcaaaaggttatggacgagacc
ctgcgctcgttagacgaaaactttaaaagcctgtcgttgggtacggcaccgttgttgttggcttgtgatttggagagtgcagccatgcaggagtactgccgtactc
tgttcaacatgcgcccggacattgcttgctgcattacgcgtatgatttgcgggctggatttgcgtccatatctgggacacgtgaccgttccatgccacatcattcaa
agctcgaacgacatcatggttccggtcgcggtgggggagtacttgcgtaagaatcttggtggcccatcggtcgttgaggtgatgcccaccgagggacatttac
cgcatttgagtatgcctgaagttactatccctgtagtgttacgtcacattcgccaagacatcacggaccac

**AS-cpGFP-AS**

gcatcgtataacgtctttatcatggccgacaagcagaagaacggcatcaaggcgaacttcaagatccgccacaacatcgaggacggcggcgtgcagctcgc
ctatcactaccagcagaacacccccatcggcgacggccccgtgctgctgcccgacaaccactacctgagcgtgcagtccaaactgagcaaagacccaacg
agaagcgcgatcacatggtcctgctggagttcgtgaccgccgccgggatcactctcggcatggacgagctgtacaagggcggtaccggagggagcatggt
gagcaagggcgaggagctgttcaccggggtggtgcccatcctggtcgagctggacggcgacgtaaacggccacaagttcagcgtgtccggcgagggcga

| |
|---|
| gggcgatgccacctacggcaagctgaccctgaagttcatctgcaccaccggcaagctgcccgtgccctggcccaccctcgtgaccaccctgacctacggcg<br>tgcagtgcttcagccgctaccccgaccacatgaagcagcacgacttcttcaagtccgccatgcccgaaggctacattcaggagcgcaccatcttcttcaagga<br>cgacggcaactataagacacgcgctgaggttaagttcgagggcgacactctggttaaccgcatcgagctgaagggcatcgacttcaaggaggacggcaaca<br>tcctgggccataagcttgaatataacttcaacgcgtcc |
| **LSSmOrange** |
| ggtggagaagcagcagcaaaggaagcggcggcaaaagggggcatggtgagcaagggcgaggagaataacatggccatcatcaaggagttcatgcgctt<br>caaggtgcgcatggagggctccgtgaacggccacgagttcgagatcgagggcgagggcgagggccgcccctacgagggctttcagaccgttaagctgaa<br>ggtgaccaagggtggcccccctgcccttcgcctgggacatcttgtcccctcagttcacctacggctccaaggcctacgtgaagcaccccgccgacatccccga<br>ctacctcaagctgtccttccccgagggcttcaagtgggagcgcgtgatgaacttcgaggacggcggcgtggtgaccgtgactcaggactcctccctgcagga<br>cggcgagttcatctacaaggtgaagctgcgcggcaccaacttcccctccgacggccccgtaatgcagaagaagaccatgggcatggaggcctcctccgagc<br>ggatgtaccccgaggacggcgccctgaagggcgaggacaagctcaggctgaagctgaaggacggcggccactacacctccgaggtcaagaccacctac<br>aaggccaagaagcccgtgcagttgcccggcgcctacatcgtcgacatcaagttggacatcacctcccacaacgaggactacaccatcgtggaacagtacga<br>acgcgccgagggccgccactccaccggcggcatggacgagctgtacaag |
| **B42AD-MAX2** |
| aaaaaataattctttcataatgggagccccgccgaaaaaaaagaggaaggtcgccgggataaataaagacatcgaggaatgcaatgccatcatcgaacaattt<br>attgactatctgagaacagggcaagagatgccaatggagatggccgaccaggctattaacgttgtaccgggcatgaccccaaaaacgatcttgcacgctggtc<br>cccgattcagcccgattggcttaaatctaacggttttcacgagatcgaggccgacgtcaatgacaccagccttctactaagtggcgatggaggaggagggtct<br>gggggcggaggcagcatggctactcagctgaatgacttgcctgacgttatacttagtaatattattgcagcggttactgacgttagaagcagaaactcaacgag<br>ctttgtctgtagaaaatggttagtgctggagcgttcaactcgtgttagccttacgcttagaggcaatgtaagagacttatttatgctgcccacatgttttaggagtata<br>acacacctagatctttcattaatttcaccctggggacatccacttctgtccccgactacgccggacccatctcttaccgctcatttactacatcacgcgtttcccttcg<br>tcacatccctagtggtatacactaggcacccccttcactcttcagttattaccgcccctatggcctcagttgaaacaaatcaagctagttaggtggcaccagaggcc<br>acaacttgcgacaggggatgagtttaacatgttattcgagaactgtccaaacttgtcaagccttgatttaagcacgttttactgttggacggatgatattcccacggc<br>cctggtctcccatccgatggttgcgagtaatttggtaaccctaaatctattaaacccctgttttagcgaggggttcaaaactgatgagattaaggctatcacgttggc<br>atgcccgaatttgaaagaatttcgtgttgtgtgcatgttcgacccgaggtacatcggttttgtgggcgacgaaggattggttgccgtcgcgacgaactgtccgaa<br>gttatccacattgcatcttgcggacaccagtgccttatctaattcaagaggggacattaatgacgatggcttcacgcaagaagatgctaagttcggagttagcacc<br>ctgatagaggtttttctggcttaccccctgctggaggagttagtactggatgtctgtaataatgtgcgtgatacgggacccgcgcttgagattttaaataaaaaatgc<br>ccaaggcttagatctctgaagcttgggcaattccatggaatttccatgcccgtggaatctaagcttgatggtgtagcgctatgccagggcctggaaagcttgagt<br>ataaggaatgtgggcgacttgaatgacatgggcctaatcgcaattggcagaggttgttctaggctagccaagtttgaagttcaaggatgtaagaaaataacagtt<br>aggggcatgcgtactctagcatcccttttgaagaagacactaattgacgttaagatctcatgttgcaaaaacttaggcgctgcgtatagcttaaaagcattagagc<br>cgattcagaataggattcagaaacttcacatagactgcgtctgggattccgtcgaagagttcgaaaatctagatggctatggatacggctttgatctgaacagaa<br>gagatggatgtgaagctagctccaatttcggggatacctttggatgcgaggaagatgcttacttgtttaaggagaagaaaaggtgtaaatttagctacgatctaaa<br>ttcactgtacgaggaggtgaatgggcatggtaacggttatagcgggcgttcctgggatagactacagtacctaagtctatggatcggagttggcgatcttctaac<br>cccgctaacggccgccgggttagaagactgccctaatttggaagaaataaaaatcagggtagagggagattgcagactatggtccaaacattcagaacaggc<br>atttggacttagcaccctattgcattaccccaaattatcaaaaatgcacttagactgtggggacacaatcgggtatgcacataccgcgccctccggtcaagtggat<br>ctgagtctatgggagaggttttacctgctagggattgggaccctttctctaacggagctagactattggccccctcaagatatggatgttaatcagaggtgtctgtc<br>tctgcccgccgcaggccttttgcaagaatgtctaactttaaggaaattgttcatacacggcacggcgcacgagcactttatgatgtttttgcttcgtatccccaatct<br>gagggatgtgcagttaagagaagattactatccggccccagaaaatgatatgtcaactgagatgcgtgccgatagcttatcaaggtttgaagcagccttaaatcg<br>tagacctatctcagactaa |

Supplementary table 3: primers used in this study

| **General primers for domain insertion biosensors** | | |
|---|---|---|
| DAD2N-F1 | attttgtttaactttaagaaggagatataccatgggtcaaactttactggacg | |
| DAD2N-F2 | catcaccatcaccatcacgagaacctgtattttcagggtcaaactttactggacgcctt | |
| DAD2N-F3 | attttgtttaactttaagaaggagatataccatgggtcatcaccatcaccatcac | |
| DAD2C-R1 | agaggccccaaggggttatgctagttattgctcagcttaacgatgagacagtgcac | |
| DAD2C-R2 | tgccgccgcttcctttgctgctgcttctccacgatgagacagtgcacg | |
| LSSmOrange-F | caagagcttaggcgtgcactgtctcatcgtggagaagcagcagcaaag | |
| LSSmOrange-R | agaggccccaaggggttatgctagttattgctcagccttgtacagctcgtccatgc | |
| **Primers conferring cpGFP insertion sites** | | |
| cpGFP insertion site | Reverse primer | Forward primer |
| DAD2 A151 | gtcggccatgataaagacgttatacgatgctgcctcca<br>tcgcggaaaa | cataagcttgaatataacttcaacgcgtccaactatgag<br>gcgtgggtgaatg |
| DAD2 G166 | gtcggccatgataaagacgttatacgatgcgcccact<br>gccaaaggcgc | cataagcttgaatataacttcaacgcgtccgctgatgtac<br>cggcagcg |

| | | |
|---|---|---|
| DAD2 A167 | gtcggccatgataaagacgttatacgatgcagcgccc actgccaaagg | cataagcttgaatataacttcaacgcgtccgatgtaccg gcagcggtaag |
| DAD2 N182 | gtcggccatgataaagacgttatacgatgcgttaaaca gtgtacgggagaattc | cataagcttgaatataacttcaacgcgtccgtaccggca gcggtaaggg |
| DAD2 M183 | gtcggccatgataaagacgttatacgatgccatgttaa acagtgtacgggaga | cataagcttgaatataacttcaacgcgtcccgtccagac ataactttgttcg |
| DAD2 R184 | gtcggccatgataaagacgttatacgatgcacgcatg ttaaacagtgtacgg | cataagcttgaatataacttcaacgcgtccccagacata actttgttcgttagtagg |
| ShHTL7 C164 | gtcggccatgataaagacgttatacgatgcacaagcc aacaacaacggt | cataagcttgaatataacttcaacgcgtccgatttggag agtgcagccat |
| ShHTL7 D165 | gtcggccatgataaagacgttatacgatgcatcacaa gccaacaacaacg | cataagcttgaatataacttcaacgcgtccttggagagt gcagccatg |
| ShHTL7 L166 | gtcggccatgataaagacgttatacgatgccaaatca caagccaacaacaa | cataagcttgaatataacttcaacgcgtccgagagtgca gccatgcag |
| ShHTL7 E167 | gtcggccatgataaagacgttatacgatgcctccaaat cacaagccaacaac | Cataagcttgaatataacttcaacgcgtccagtgcagc catgcaggagt |
| **Primers for Y2H-based biosensor and protoplast plasmid construction** | | |
| 32 | aacctccttggattagtacatgcgtagtgagttcggtgtt | |
| 33 | aacaccgaactcactacgcatgtactaatccaaggaggtttacg | |
| 101 | caagatatttcttgaatcaggcatttaaatttaaggccacataaatcc | |
| 102 | tttttgaaaattcaatataacactgtgctttctgttgctg | |
| 103 | cagcaacagaaagcacagtgttatattgaattttcaaaaattcttacttt | |
| 104 | ctagccgtcagagctttcattatagttttttctccttgacgttaaag | |
| 105 | gtcaaggagaaaaaactataatgaaagctctgacggctag | |
| 106 | attgggctacgtaaattcgattaacgatgagacagtgcac | |
| 107 | tgcactgtctcatcgttaatcgaatttacgtagcccaatc | |
| 108 | tttttcggcggggctcccattatgaaagaattatttttttattatgttaatct | |
| 109 | aaaaaaaataattctttcataatgggag | |
| 110 | ctcttttgtttttctcgagattagtctgagataggtctacgatttaag | |
| 111 | gtagacctatctcagactaatctcgagaaaaacaaaagagttaat | |
| 112 | tttttgaaaattcaatataattggtcgtttgtcatcaag | |
| 113 | cttgatgacaaacgaccaatt | |
| 117 | gtcaaggagaaaaaactataatgtctaaaggtgaagaattattcac | |
| 118 | acgatttgctctcaatccgcttaagatcttttgtacaattcatcc | |
| 120 | aattgtacaaaagatcttaagcggattgagagcaaatc | |
| 121 | agattccggccatatttctgacgccacagaaacctcag | |
| 122 | ttctgaggtttctgtggcgtcagaaatatggccggaatc | |
| 123 | aaagggaacaaaagctggagctatttaaatggtagcttttcgttcc | |
| 124 | aattcttcacctttagacattatagttttttctccttgacgttaaag | |
| 183 | gttatggtgttcaatgttttgc | |
| 184 | tgcatcaaaaaaaaaaattatccgatgatgagcacgccacagaaacctcag | |
| 185 | gctcatcatcggataatttttttttttgatgcatctatacagagtgcaccataccacag | |
| 186 | agattccggccatatttctgcaccgcatagggtaataactg | |
| 187 | agttattaccctatgcggtgcagaaatatggccggaatc | |
| 188 | caatttcacacaggaaacagc | |
| 441 | taactaatcgatatgggtcaaactttactggacg | |
| 442 | aatattactagtttattgctcagccttgtacag | |
